# MMTF - an efficient file format for the transmission, visualization, and analysis of macromolecular structures

**DOI:** 10.1101/122689

**Authors:** Anthony R. Bradley, Alexander S. Rose, Antonín Pavelka, Yana Valasatava, Jose M. Duarte, Andreas Prlić, Peter W. Rose

## Abstract

Recent advances in experimental techniques have led to a rapid growth in complexity, size, and number of macromolecular structures that are made available through the Protein Data Bank. This creates a challenge for macromolecular visualization and analysis. Macromolecular structure files, such as PDB or PDBx/mmCIF files can be slow to transfer, parse, and hard to incorporate into third-party software tools. Here, we present a new binary and compressed data representation, the MacroMolecular Transmission Format, MMTF, as well as software implementations in several languages that have been developed around it, which address these issues. We describe the new format and its APIs and demonstrate that it is several times faster to parse, and about a quarter of the file size of the current standard format, PDBx/mmCIF. As a consequence of the new data representation, it is now possible to visualize structures with millions of atoms in a web browser, keep the whole PDB archive in memory or parse it within few minutes on average computers, which opens up a new way of thinking how to implement efficient algorithms in structural bioinformatics. The PDB archive is available in MMTF file format through web services and data are updated on a weekly basis.

## Introduction

The Protein Data Bank (PDB) [1] is the global archive of 3D structures of proteins, nucleic acids, and complex assemblies. Recent advances in experimental techniques have led to an explosion in both the number and size of such structures. The entire PDB now exceeds one billion atoms and the largest structure currently contains about 2.4 million atoms [2] (Fig 1A). In addition to a growing number of depositions per year (Fig 1B) and average number of atoms per structure (Fig 1C), 68 of the 100 largest structures were deposited in the past three years. In Fig 1D, we show the rising importance of Cryo-Electron microscopy as a technique [3]. It is expected that much larger molecular machines and molecular assemblies will be modeled by combining multiple experimental techniques [4].

**Fig 1.**
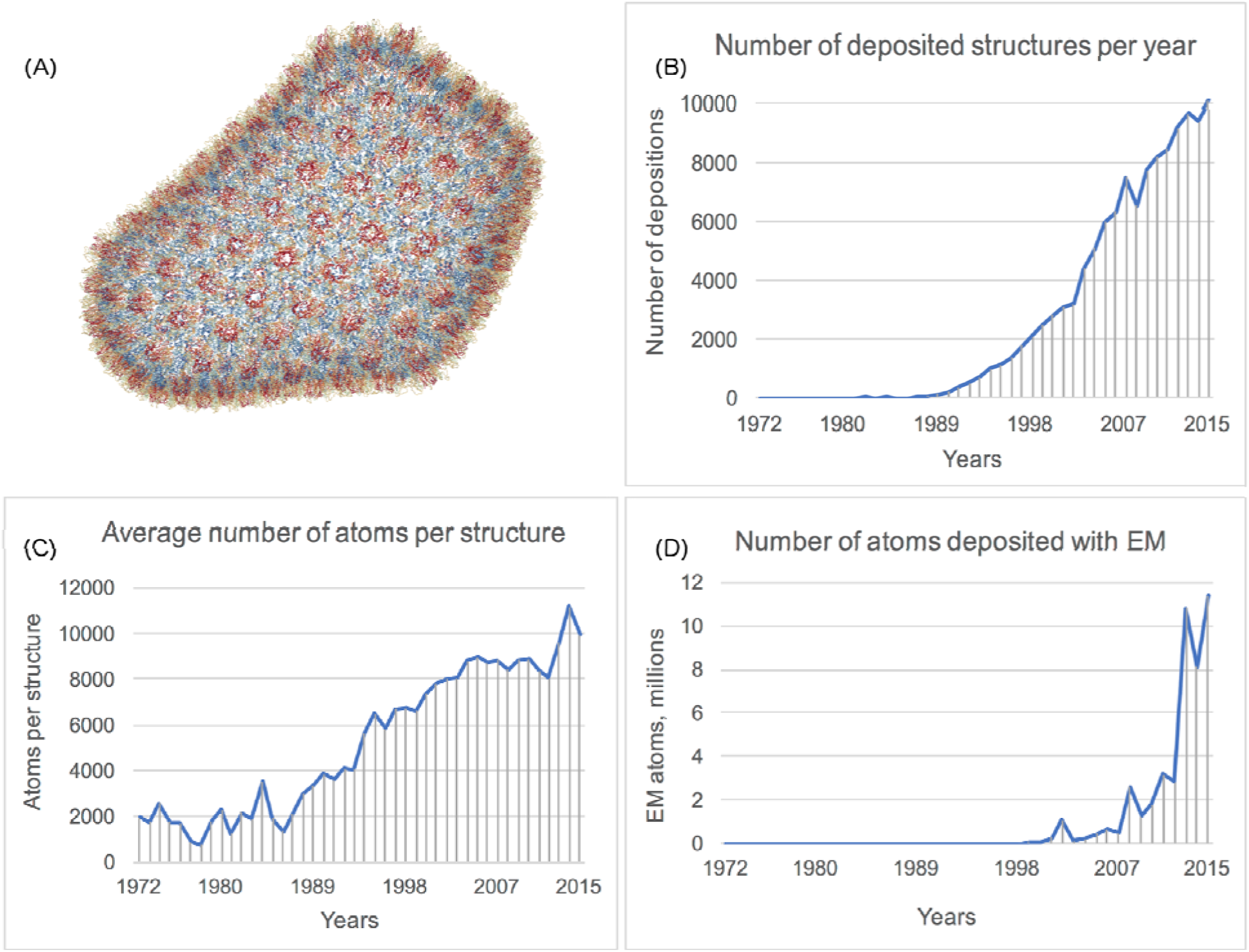
Growth of the Protein Data Bank Archive. (A) The currently largest asymmetric structure in the PDB - the HIV Capsid (PDB ID 3J3Q) contains over 2.4 million atoms. (B) The number of depositions per year (obsoleted or superseded entries are excluded). (C) The average structure size (asymmetric unit size for crystallographic structures). (D) Electron microscopy structures are contributing ∼10 million atoms per year for the past 3 years (1% of the archive).

Significant increases in data sizes have been seen in many fields. Efficient storage and transmission of data using novel file formats and data compression methods are integral to these development, e.g., for the transport of HD-TV, video, and audio. A similar trend has emerged in the handling of whole genome data [5].

Two notable developments have been made in developing such a format for macromolecules. First, WPDB [6] stored the data as binary files with limited precision, allowing efficient access. WPDB is however no longer maintained and was tied to the Windows operating system. The second development is mmJSON [7], which represents data from the PDBx/mmCIF format (http://mmcif.wwpdb.org/) in the JSON serialization format that can be efficiently parsed by modern web browsers. Even after compression with gzip (a commonly used general purpose compression tool) the largest structure (PDB ID 3J3Q) takes up 27 MB of space making it challenging to transfer - in particular over mobile networks. Moreover, neither WPDB, mmJSON, nor other formats such as PDBx/mmCIF, provide all data necessary to represent a full macromolecular model including bond information. Furthermore, as text based formats, they are slow to parse, and clean Application Programming Interfaces (APIs) are generally not made available.

Commercial software providers have produced their own internal representations of macromolecular structures. No such format, however, is openly available and thus they cannot be incorporated into third party software or developed with community involvement. For this reason, structural analysis is currently a laborious and error-prone process, often involving substantial duplicated effort to reliably process the entire PDB archive into a 3rd party data structure. Structure visualization can be equally challenging for large structures, due to slow data download and high client-side memory requirements to parse large structure files. Some of the largest structures in the PDB require more memory than is typically available within in web browser.

In this paper we describe a new data representation, the **M**acro**M**olecular **T**ransmission **F**ormat (MMTF) (http://mmtf.rcsb.org/) that aims to resolve these deficiencies. MMTF is a binary machine-readable file format that can be parsed, in some instances at least an order of magnitude faster than existing text-based formats. Custom lossless and lossy compression methods with either full atom level detail and a reduced representation (C-alpha, P atoms) are applied [8] to reduce the file size and thus further improve transmission and parsing speeds. Finally, MMTF is designed for interoperability and use by a broad community. APIs are provided in common programming languages and a full chemical description required to understand a structure is included in the file. The PDB archive is provided in MMTF format through web services and updated weekly. MMTF is a macromolecular file format for the modern age.

## Design and Implementation

### Design Considerations

Above we demonstrated that existing file formats are becoming less suitable for modern macromolecular data. Due to these challenges, the MMTF format was designed with three core aims. First, to minimize data storage requirements and transfer times, the format should represent data in compressed form without loss of accuracy. Second, it should be fast to parse, since I/O is often a bottleneck in structural analysis and visualization. Third, we designed MMTF to be as extensible, self-contained, and interoperable as possible. As a binary, machine-readable format, the preferred access to MMTF data is through the APIs provided in several programming languages. This allows the developers to focus on scientific applications and not on developing file parsers.

### Data Items and Encoding

The MMTF format was designed to include the core data commonly used by macromolecular visualization and analysis tools (Table 1). A comprehensive list of the data items is available in the MMTF specification. Additional data, if required, can be accessed through web services.

**Table 1.** Data Categories described in MMTF format.

| Data Category | Data Items |
| --- | --- |
| Metadata | PDB ID, title, deposition date, release date, experimental method(s) |
| Crystallographic info | Space group, unit cell, NCS operators, resolution, $R_{\text{free}}$, $R_{\text{work}}$ |
| Primary structure | Polymer sequences |
| Secondary structure | DSSP secondary structure assignments* |
| Structural model | Models, chains, groups (residues), atoms, bonds* and bond orders* |
| Quaternary structure | Biological assembly transformations |

\* These data items are not available in the PDBx/mmCIF files and are added to MMTF files.

To make MMTF files directly usable by visualization and analysis applications, we add consistently calculated DSSP [9] secondary structure information using the BioJava implementation [10]. MMTF includes the full chemical description of all molecules in a PDB entry. Bonds and bond orders for both standard and non-standard residues, e.g., ligands, are included from the Chemical Component Dictionary [11] and additional covalent bonds (struct_conn category in the PDBx/mmCIF files), such as disulfide bonds or covalent bonds between ligands and polymers are also included in MMTF. Metal coordination and hydrogen bond information is not included in MMTF, since there are no generally agreed upon standards how to define them. Fig 2 describes the creation of an MMTF file from a PDBx/mmCIF archive file.

**Fig 2.**
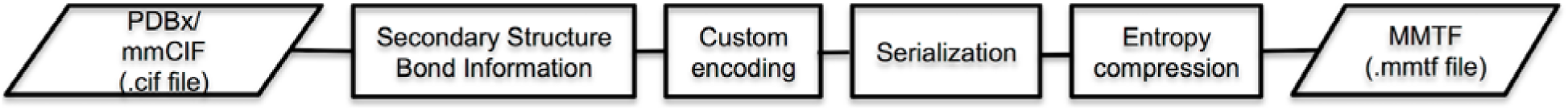
Steps in the creation of a MMTF file from a PDBx/mmCIF file. After parsing a PDBx/mmCIF file, DSSP secondary structure is calculated and bond information is added for all residues. Custom encoding strategies are applied to the different data categories to achieve a compact representation. These data are serialized in binary form and then further compressed with standard compression tools to create a compressed MMTF file.

### Encoding Strategies

In order to reduce the overall file size, we applied specialized encoding techniques to make the data more compressible. These techniques either reduce redundancy in the data or reduce the dynamic range (entropy) of numbers, to make them more compressible using standard entropy encoding techniques.

Fields of the same type are grouped together in MMTF to create a flat data structure. For instance, the coordinates of all atoms are stored together, instead of in atom objects with other atom-related data. This avoids imposing a deeply nested hierarchical structure on consuming programs, while still allowing efficient traversal of models, chains, groups, and atoms. This approach represents a columnar encoding [12] of data, which facilitates data encoding and enhances data compressibility.

Lossless integer encoding is applied to all fixed precision floating point numbers. Integer numbers have a simpler bitwise representation and are therefore more compressible than the equivalent floating-point numbers [8]. Atomic coordinates are typically represented with a precision of 3 decimal places, and temperature factors with 2 decimal places. For lossless encoding, we multiply coordinate and temperature factor values by 1000 and 100, respectively, and round the values to the nearest integer.

A further increase in compression can be obtained through lossy encoding by rounding coordinates to 0.1 Å precision and temperature factors and occupancy to 0.1 precision. Lossy compression is particularly important for the visualization of large complexes, for which the reduced precision is not visually perceptible [6,8].

Dictionary encoding is used for data repeated across multiple residues. In standard PDB and mmCIF files, atoms within a residue are listed in a standard order. Exploiting this, atom name, element symbol, intra residue bonds and bond orders, etc. can be stored once for each unique residue type and not repeated across the file, as shown for the dictionary entry for serine (Fig 3). MMTF has been designed to handle exceptions to a consistent atom order, if they occur, however, the encoding will be less efficient.

**Fig 3.**
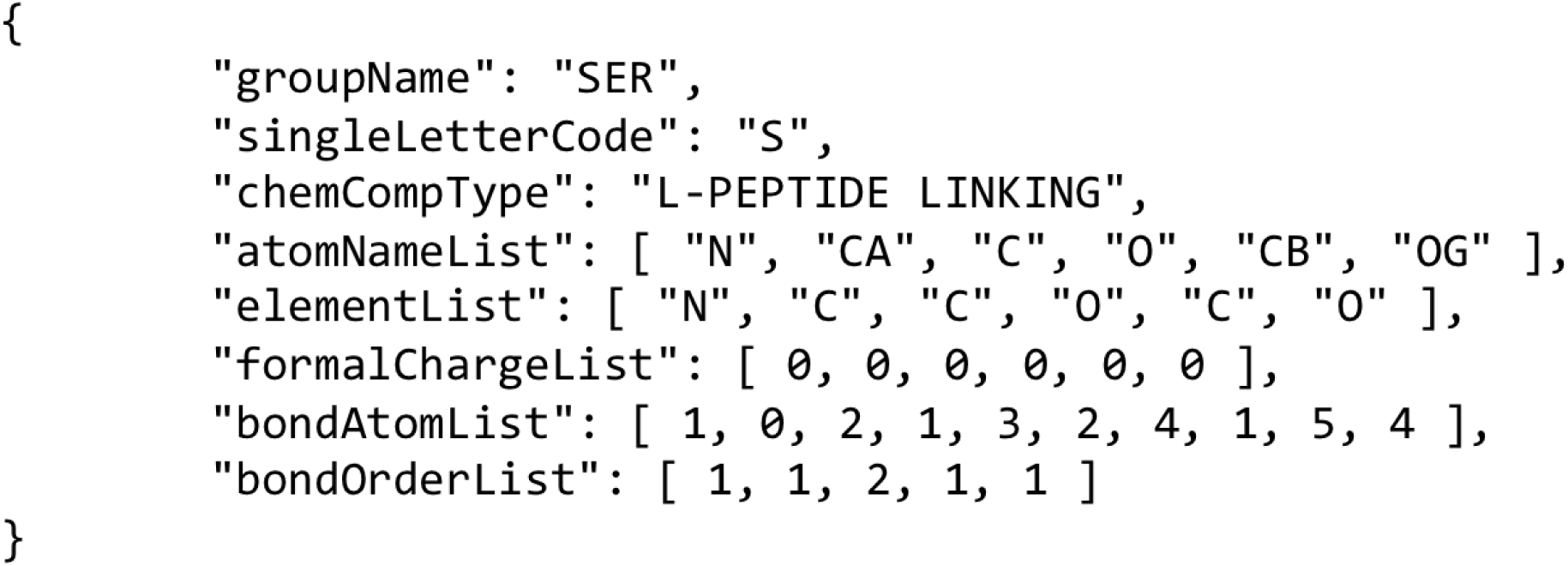
Dictionary entry for amino acid serine.

Delta encoding is applied to data of large magnitude that change in small increments. For example, instead of storing absolute atom coordinate values, differences in the x, y, and z coordinates are stored. Due to the covalent bond structure in molecules, these differences typically lie within a small dynamic range bound by their bond distances. Previous work determined this method to be the most effective encoding technique [8]. Temperature factors are also delta encoded, since their variation from residue to residue is typically low.

Run-length encoding compresses a list of repeated values, such as occupancy values in X-ray structures, most of which are constant (1.0). Here the value itself and the number of repetitions is stored. For atom serial numbers, delta and run-length encoding are combined to achieve a very compact encoding.

Recursive indexing - Given the small dynamic range of delta encoded coordinates, most, but not all values can be represented as 16-bit signed integers, rather than 32-bit signed integers. We have explored the effect of packing on compression [8] and identified recursive indexing as a simple and effective packing strategy for this data type. Any values that lie outside the 16-bit integer range [-32,768, 32,767] are decomposed into a series of values, such that the individual values fit into the 16-bit range (Fig. 4D), and their sum adds up to the original value.

The overall workflow for the encoding of columnar data is shown in Fig 4.

**Fig 4.**
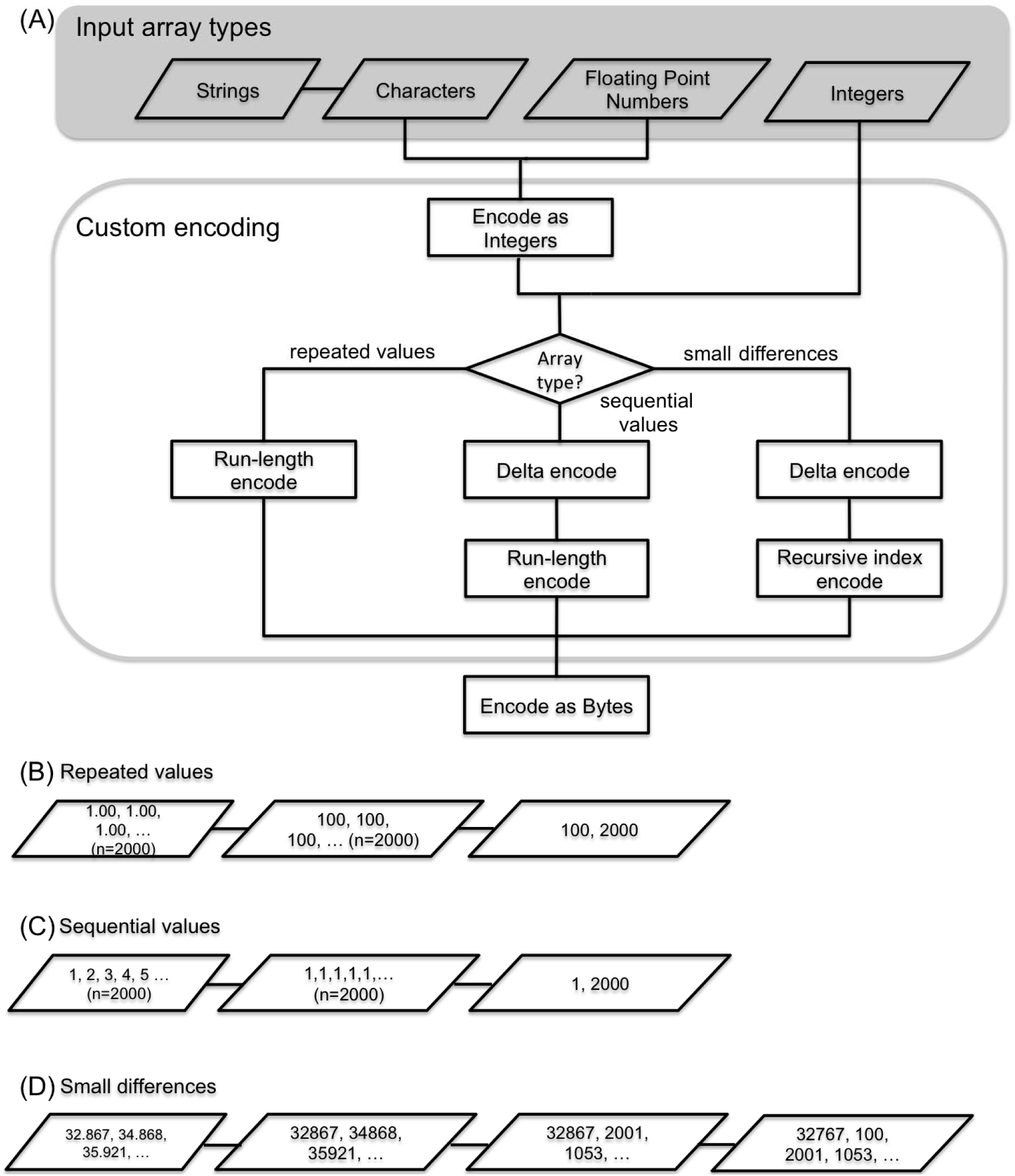
Workflow for encoding columnar data within MMTF. (A) Columnar data are first converted to integer arrays. Depending on the type of the values in the array, three types of custom encoding are applied to: 1. Repeated values, 2. Sequential values, 3. Small differences between adjacent values. All encoded values are finally encoded as a byte array. (B) Example of encoding 2,000 occupancy values by integer encoding (x100) followed by run-length encoding. (C) Example of encoding 2000 atom serial numbers by applying delta and run-length encoding. (D) Example of encoding atom coordinate values by integer encoding (x1,000), delta encoding, and recursive index encoding into a 16 bit signed integer array. Here, the value 32,867 exceeds the maximum value (32,767) for a 16-bit signed integer. Therefore, recursive index encoding decomposes this value into two numbers 32,767 and 100 that sum up to the original value. All subsequent values are within range and are represented directly by their values 2,001, and 1,053.

### Serialization

MMTF data are stored in the MessagePack format (version 5, http://msgpack.org) binary container format. MessagePack is an efficient binary serialization format, similar to JSON, but faster to parse and more compact. Encoding and decoding libraries for MessagePack are available in many languages. The top-level of the container holds the field names as keys and field data as values. Non-columnar data are described using standard MessagePack data types. Columnar data, e.g., most data columns in the “ATOM” records, are custom encoded. The MMTF specification defines Codec Types used to custom encode columnar data. These data records are described by the following data structure (Fig 5), which is represented as a binary array in MessagePack.

**Fig 5.**
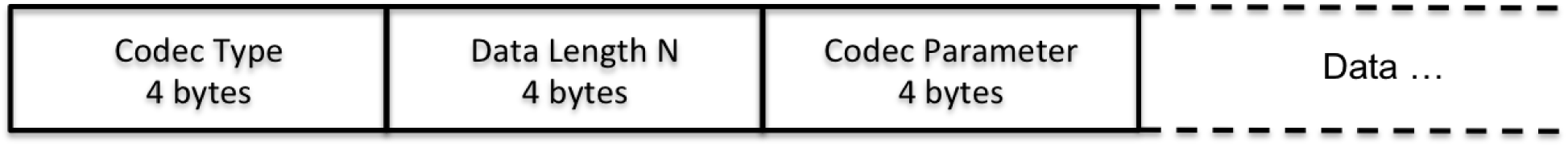
Data structure of custom encoded record in MMTF. A Codec Type describes the columnar encoding strategy. A Codec may describe the combination of several encoding strategies. For example, coordinate data are encoded by a Codec that combines integer encoding, delta encoding, recursive index encoding. Data Length represents the number of values that have been encoded, and here the Codec Parameter for coordinate encoding is a divisor to convert integers to floating point numbers.

### MMTF Data Files

MMTF files for all PDB entries are updated weekly as part of the RCSB PDB weekly update pipeline. Semantic versioning (http://semver.org) is employed to the file specification and the APIs. Major version changes of the specification may require code updates to decode and parse data. For this reason, after the release of a new major version of the specification, the previous major version will be retained for a number of months to allow time for code updates and testing. Such version changes will be disseminated through a mailing list and updates to the documentation.

MMTF files are generated with two types of molecular representation (Table 2). The reduced representation, which uses lossy compression and less atomic level detail is suitable for 3D visualization, e.g., ribbon diagrams, or calculations that require only a C-alpha representation.

**Table 2.**
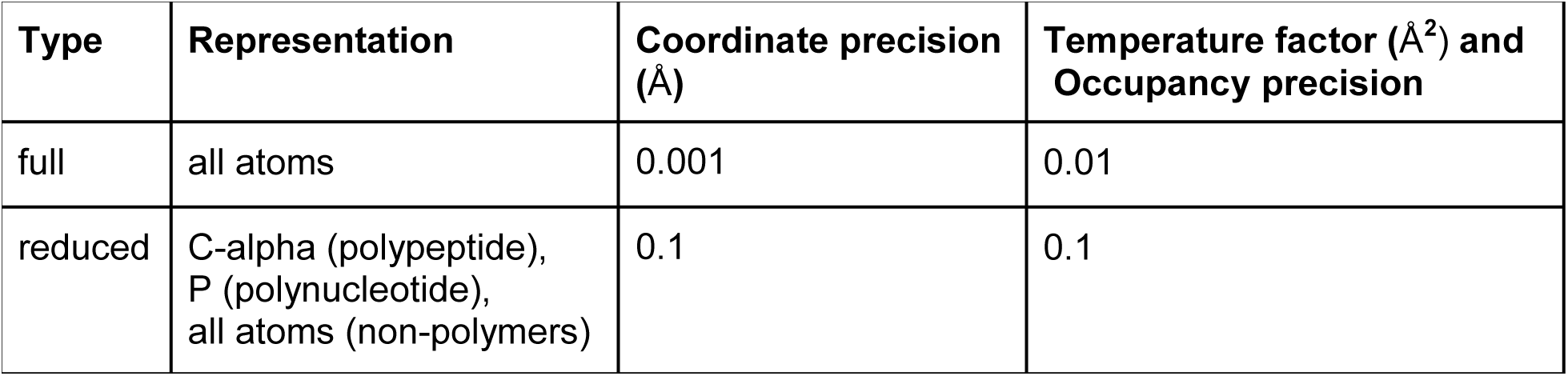
MMTF File Types.

| Type | Representation | Coordinate precision (Å) | Temperature factor (Å<sup>2</sup>) and Occupancy precision |
| --- | --- | --- | --- |
| full | all atoms | 0.001 | 0.01 |
| reduced | C-alpha (polypeptide),<br>P (polynucleotide),<br>all atoms (non-polymers) | 0.1 | 0.1 |

### MMTF Application Programming Interface

MMTF files are accessible through RESTful web services via HTTP and HTTPS protocols, or downloadable as individual gzipped files (http://mmtf.rcsb.org/download.html). A weekly update procedure ensures the availability of the latest structures, as provided by the wwPDB. For large-scale analysis of the PDB archive, where loading of thousands of individual files is inefficient, a single Hadoop Sequence file (https://wiki.apache.org/hadoop/SequenceFile) is provided. These files can be efficiently processed in parallel by Big Data frameworks such as Apache Hadoop (http://hadoop.apache.org/) or Apache Spark (http://spark.apache.org/).

The preferred access to MMTF data is via the provided decoder APIs, which are available through open source GitHub repositories. API documentation and example code are available from the MMTF project page (http://mmtf.rcsb.org/). Fig 6 shows the integration of third-party applications and software libraries with the MMTF APIs.

**Fig 6.**
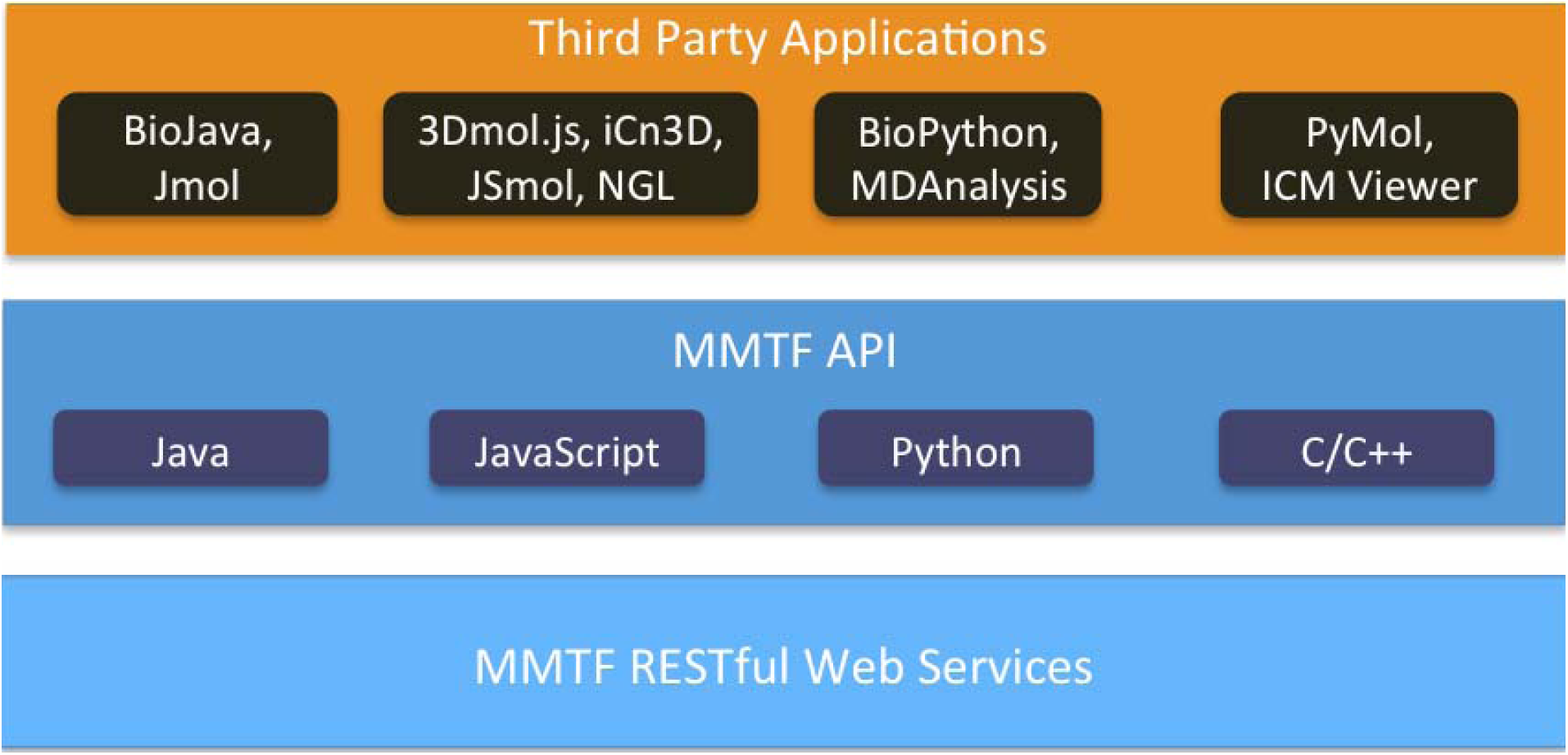
Third party software integration through MMTF APIs and web services.

MMTF data are accessed through the MMTF RESTful web services. APIs available in common programming languages provide efficient access to the MMTF data. Third party applications then access the data through the language-specific APIs.

## Results

The benefits of the MMTF file format were assessed in three different ways. First, the relative sizes of the files in different formats were measured. Second, the file load time was benchmarked in Python, JavaScript, and Java. Third, the simplicity of using the new format is demonstrated.

### File Size Comparison

In Fig 7 we compare the size of the PDB archive in mmCIF, PDB, and MMTF file formats. In the MMTF file format the PDB archive can be stored in about 8 GB, making it less than 1/4 the size of the mmCIF files and 1/3 the size of the PDB files. In practice, being stored in about 8 GB also means the entire archive can be stored in RAM on many standard desktop and laptop computers.

**Fig 7.**
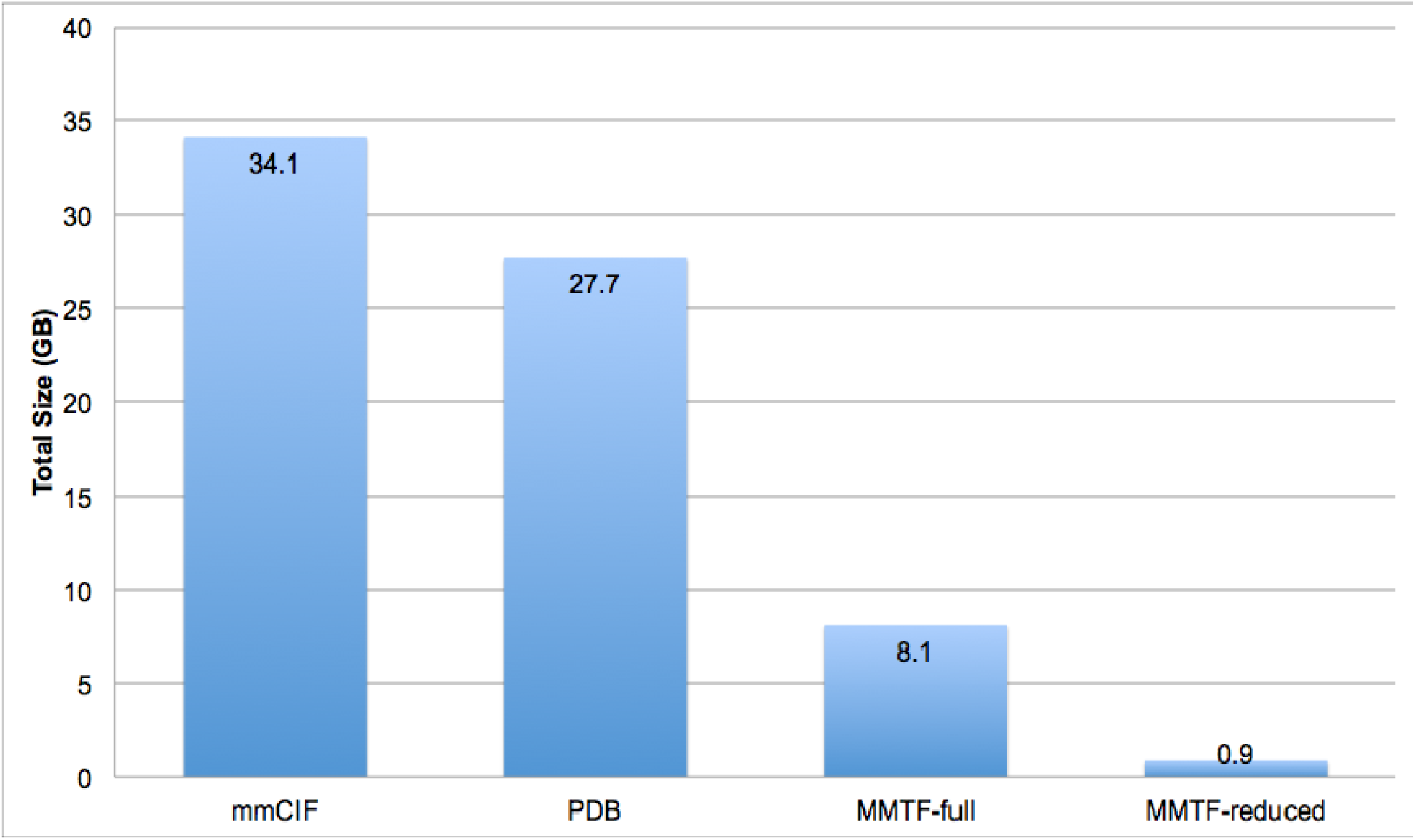
Comparison of file sizes for the PDB archive (∼127,000 entries) in gzip compressed mmCIF, PDB, and MMTF formats as of March 2017. About 500 large structures (> 99,999 atoms or > 62 chains) cannot be represented in the PDB format, however, they are available as split PDB files (.tar.gz files) and take up about 2.7 GB, which is included in the reported PDB file size. For MMTF, we report the size of the all atom representation (MMTF-full) and the reduced representation (MMTF-reduced).

### Load Time Benchmarks

The following benchmarks assess the file load time for MMTF compared to mmCIF and PDB data formats. The load times reported in the figures below consist of reading the files from a local disk, decompressing and parsing the data, instantiating a hierarchical molecular data structure (model->chain->residue->atom), and storing the metadata. All parsing benchmarks were performed using a single core on a MacMini, 2.6 GHz Intel Core i5, 16 GB RAM 1600 MHz DDR3, with a solid state drive.

The first benchmark uses the existing file parsers (mmCIF, PDB) in BioJava and compares their performance with the new BioJava MMTF parser, which uses the MMTF-Java API. In Fig 8 we compare the load times for ∼127,000 PDB entries as individual gzip compressed mmCIF, PDB, and MMTF files, and as uncompressed Hadoop Sequence files.

**Fig 8.**
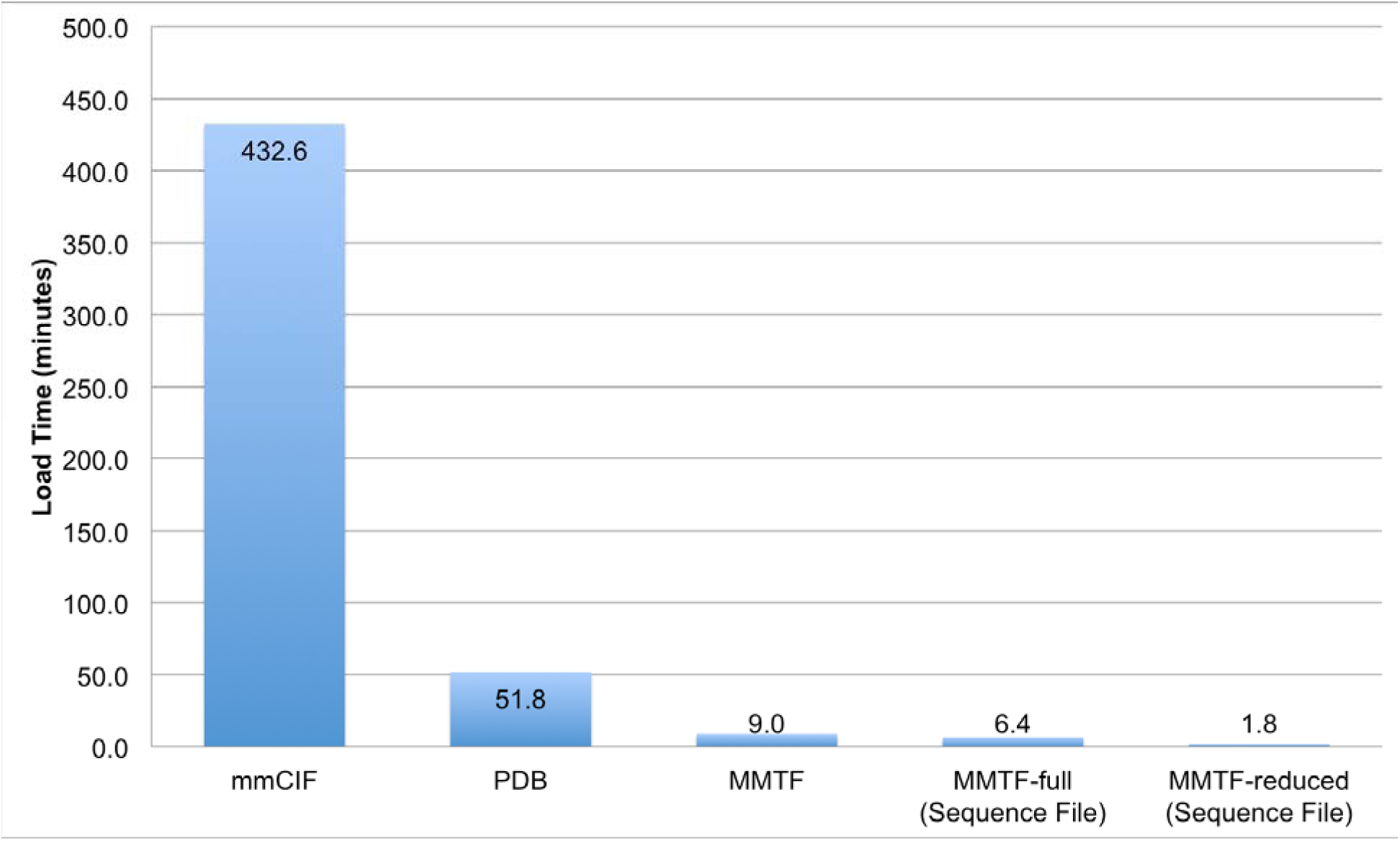
Comparison of BioJava load time for the PDB archive using different file formats. Load time for the PDB archive (∼127,000) entries using the gzip compressed mmCIF, PDB, and MMTF formats. For MMTF, we report the load time for individual gzipped files, as well as, the load time for uncompressed Hadoop Sequence Files containing MMTF records in the full (all atom, MMTF-full) and the reduced format (MMTF-reduced). For PDB file loading, about 500 large structures that cannot be represented in the PDB format (>99,999 atom, > 62 chains) were excluded.

Next, we compared the load time of implementations in different programming languages. We benchmarked three commonly used software libraries: BioPython [13] (http://biopython.org/), NGL Viewer [14] (https://github.com/arose/ngl), and BioJava [15] (http://biojava.org/) written in Python, JavaScript, and Java, respectively. A benchmark set of 1,000 randomly selected PDB entries was used for the assessment (S1 Table) (Fig 9).

**Fig 9.**
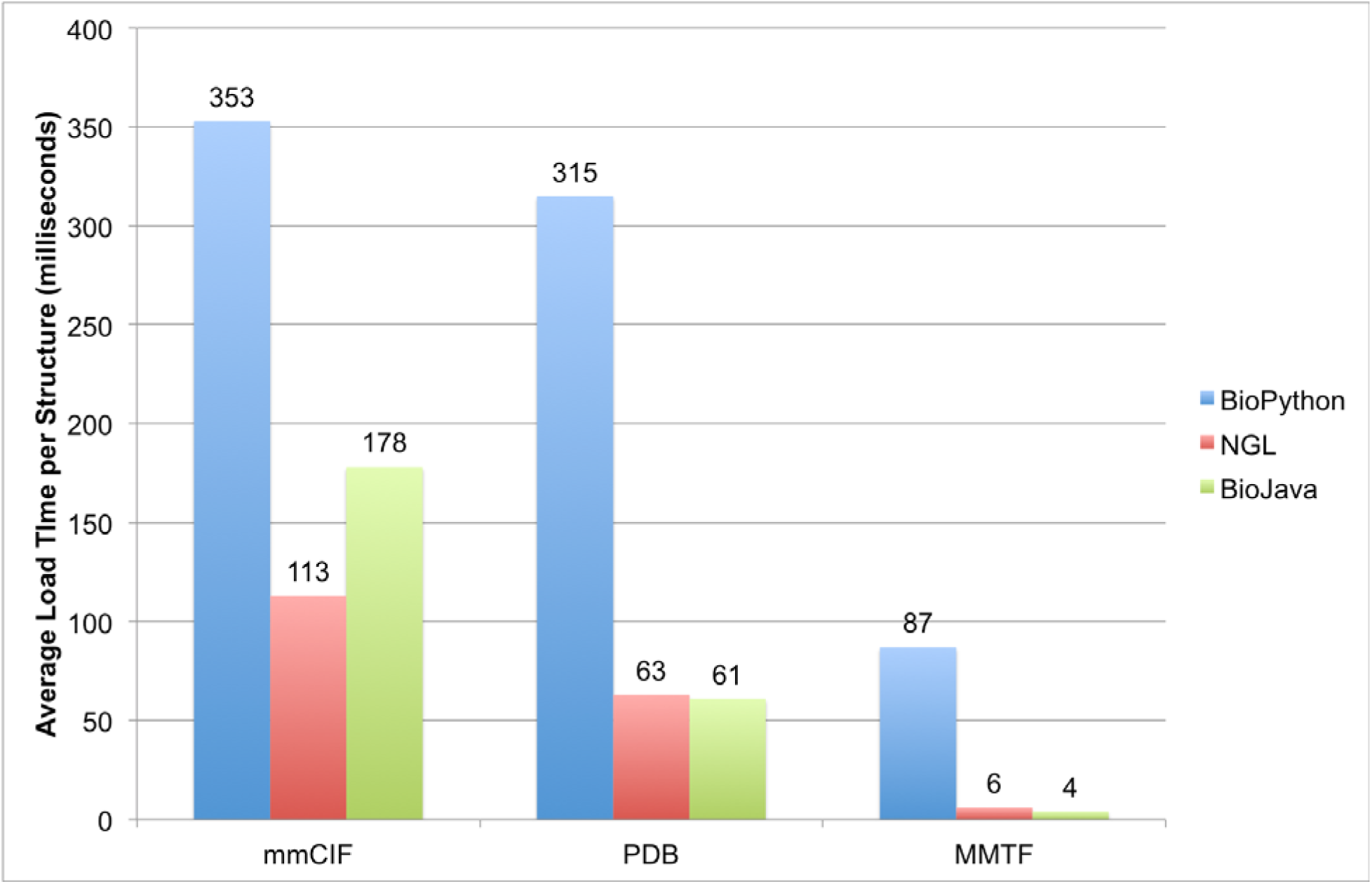
Comparison of the average load times for different file formats using three software libraries in three programming languages on a set of 1000 random PDB entries.

The MMTF format has clear advantages over mmCIF and PDB. For BioPython, MMTF is parsed about 4 times faster than mmCIF using the FastMMCIFParser (353 seconds), and about 30 times faster compared to the default MMCIFParser (2650 seconds, data not shown in Fig 9), which creates a more complete data model. For NGL (JavaScript), MMTF loading is about 18 times faster than mmCIF. For BioJava, loading MMTF files is about 45 times faster than loading the corresponding mmCIF files.

To assess the effect of structure size on load time (Fig 10), we created samples of 100 structures around the 25 percentile (S2 Table), 50 percentile (S3 Table), and 75 percentile (S4 Table) from the atom size distribution of the PDB archive. To create these subsets, we selected 100 structures symmetrically around the quartile values.

**Fig 10.**
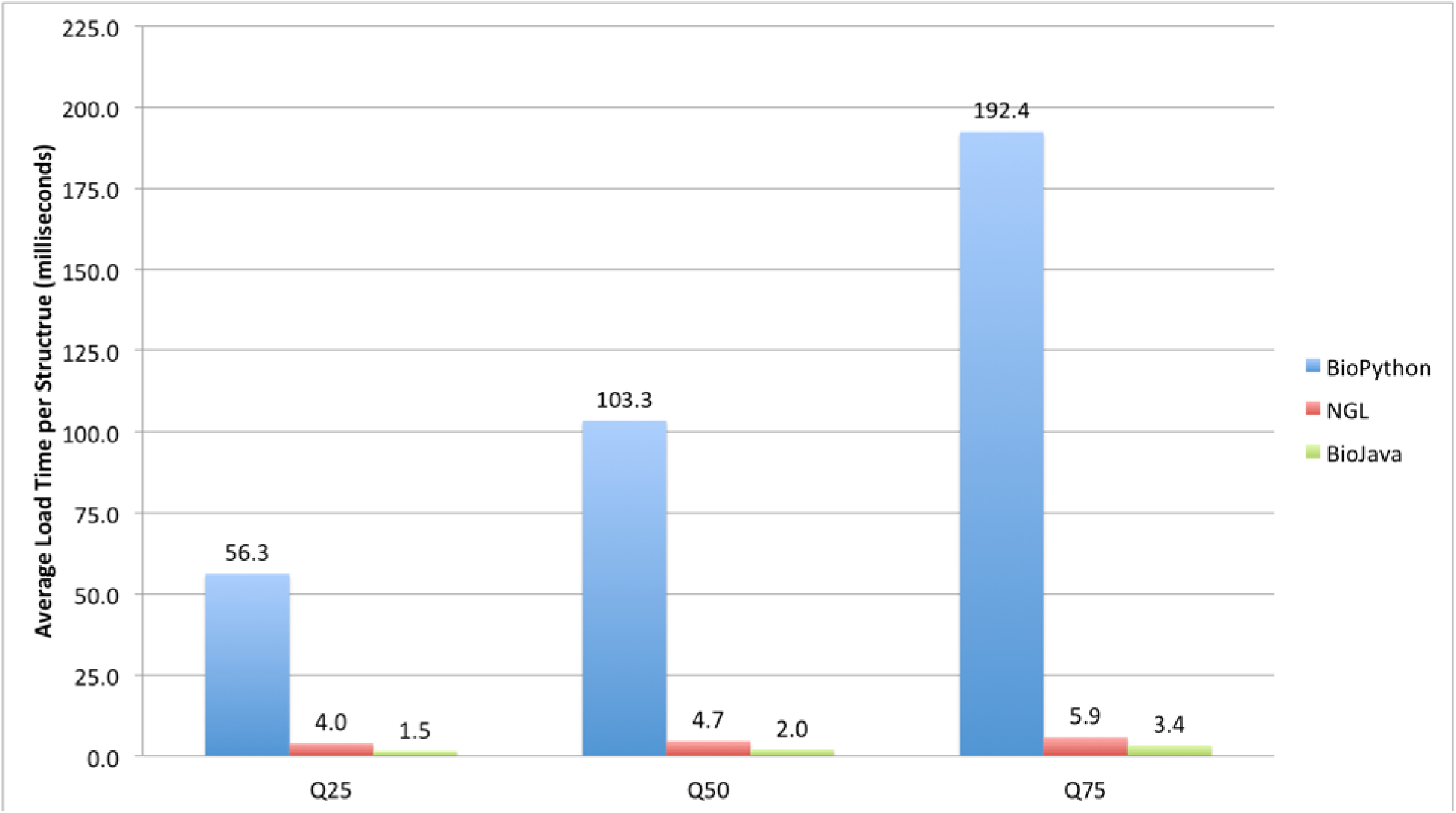
Comparison of the average load times per structure using the MMTF format for three structure sizes. The benchmarks contain 100 structures each around the 25, 50 and 75 percentile of the PDB size distribution: Q25 (2,309-2,313 atoms), Q50 (4,054-4,063 atoms), Q75 (7,862-7,885 atoms).

According to this benchmark, most small to medium sized PDB structures can be parsed in milliseconds using the BioJava/MMTF-Java API and NGL/MMTF-JavaScript API. The load time is approximately linear with the number of atoms in a PDB entry. MMTF file loading with BioPython is consistently about a factor of 40-50 slower than with BioJava. This is due in part that Python is an interpreted language. Our profiling points to the creation of the hierarchical molecular data structure as the time limiting factor for BioPython.

MMTF was specifically designed to handle the efficient transfer and visualization of very large structures that could not be parsed and visualized using the PDBx/mmCIF format due to the large memory overhead. For example, the currently largest asymmetric structure (PDB ID 3J3Q) in the PDB with 2,440,800 atoms, shown in Fig 1A, was rendered with NGL viewer using the MMTF-reduced format. Table 3 compares the load times for this entry using BioPython, NGL, and BioJava.

**Table 3.**
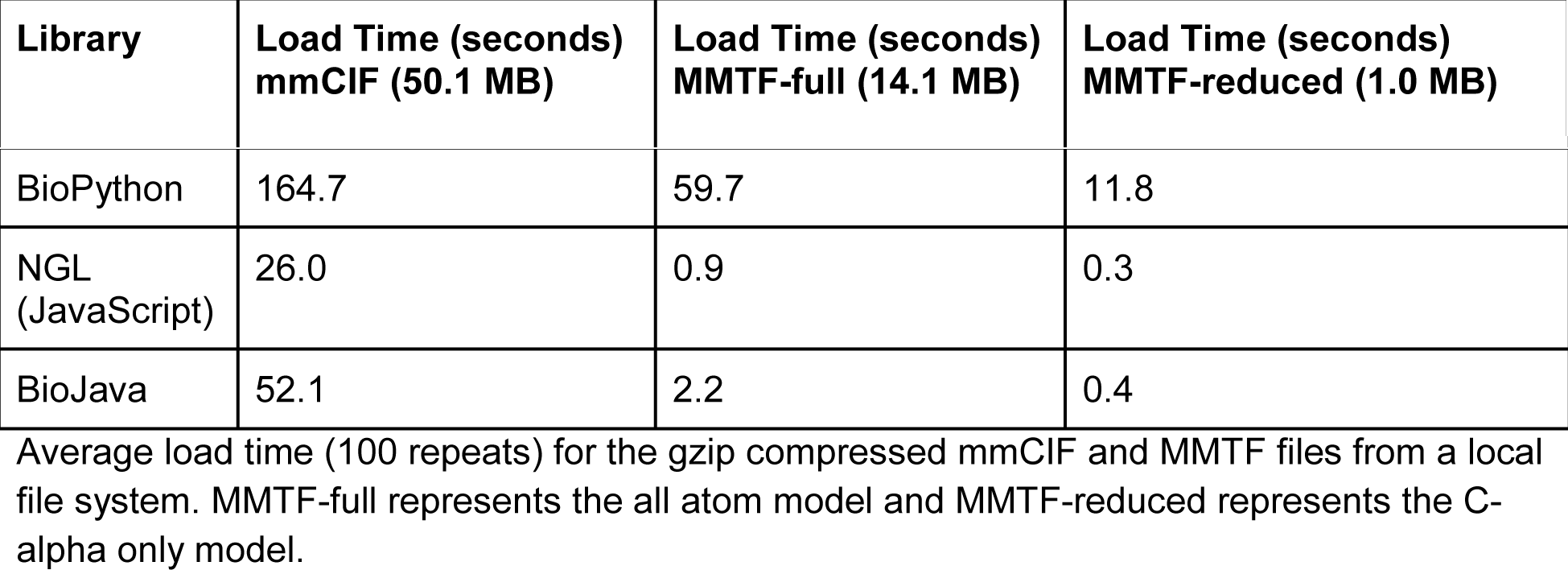
Average load time for large PDB entry 3J3Q with about 2.4 million atoms.

| Library | Load Time (seconds)<br>mmCIF (50.1 MB) | Load Time (seconds)<br>MMTF-full (14.1 MB) | Load Time (seconds)<br>MMTF-reduced (1.0 MB) |
| --- | --- | --- | --- |
| BioPython | 164.7 | 59.7 | 11.8 |
| NGL<br>(JavaScript) | 26.0 | 0.9 | 0.3 |
| BioJava | 52.1 | 2.2 | 0.4 |

Average load time (100 repeats) for the gzip compressed mmCIF and MMTF files from a local file system. MMTF-full represents the all atom model and MMTF-reduced represents the C-alpha only model.

### Simple Application Programming Interfaces

The MMTF file format is designed to be easy to use and incorporate into 3rd party applications. While MMTF has a flat, columnar data structure, it can be traversed following the structure hierarchy: Models -> Chains -> Groups -> Atoms. In Fig 11, we demonstrate the simplicity in retrieving data using the Java, JavaScript and Python APIs.

**Fig 11.**
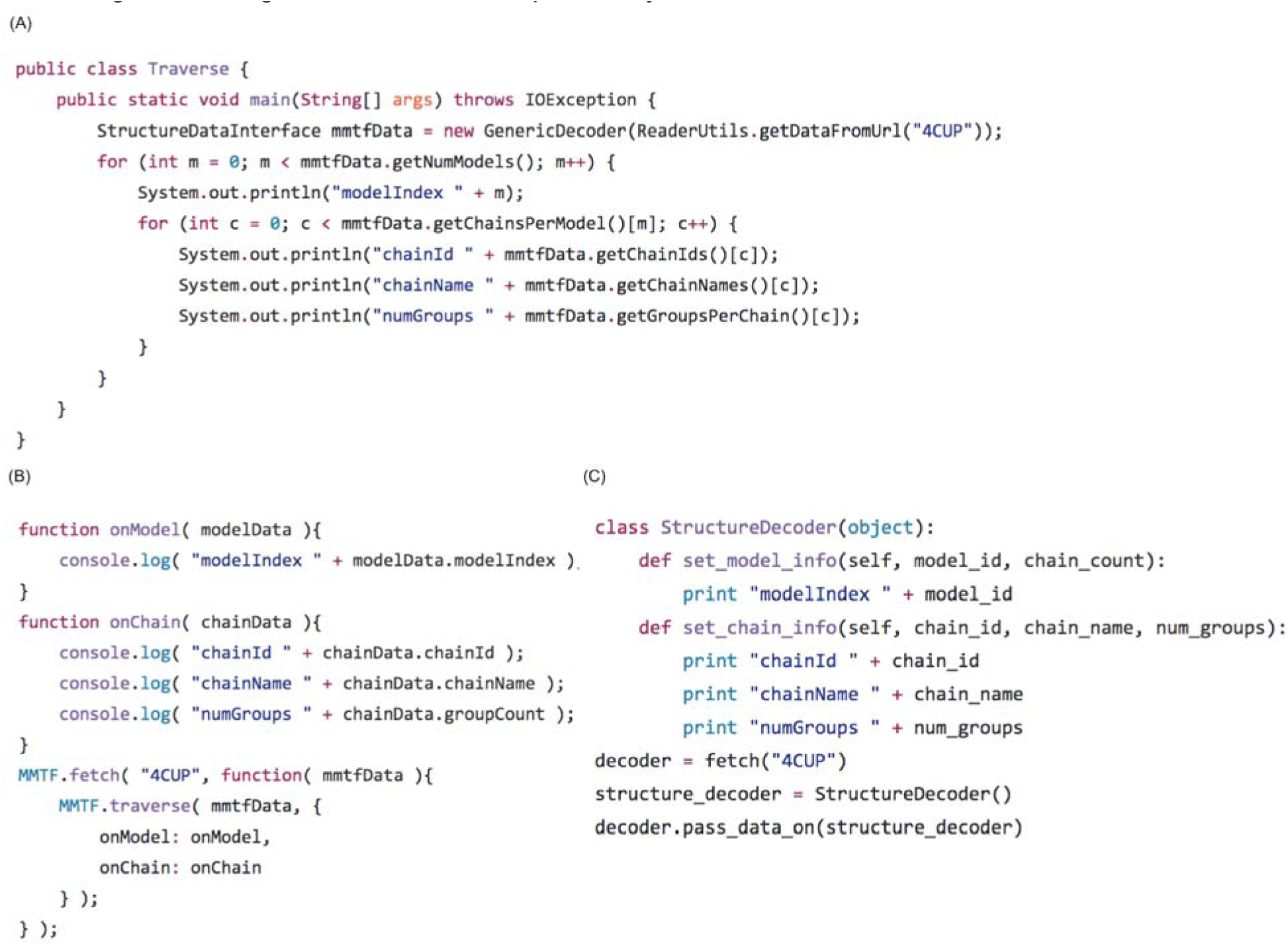
Traversal of the structure hierarchy using the MMTF API. These code snippets (A) Java, (B) JavaScript, and (C) Python demonstrate how to load and decode an MMTF file (PDB ID 4CUP) from http://mmtf.rcsb.org and then traverse the hierarchical data structure (Models -> Chains -> Groups -> Atoms). The code shown here loops through the Model and Chain hierarchy. For each model, the model index is printed, and for each chain, the chainId, chainName, and number of groups (residues) are printed. In an analogous fashion, not shown for brevity, the group and atom data can be traversed.

## Availability and Future Directions

In this paper we present a modern macromolecular transmission format. MMTF addresses the growing size and complexity of macromolecular structures through a new binary, custom compressed file format. Furthermore, MMTF is self-contained and simple APIs are provided in multiple popular programming languages. Software developers do not need to implement their own parsers - often an error-prone process, but rather build on the tools provided by MMTF. Through both these advances MMTF allows rapid user-friendly access to any structure in the PDB archive with a few lines of code. We demonstrate that the format is 75% smaller, an order of magnitude faster to parse, and is provided along with a user friendly API that promotes interoperability.

The MMTF project page (http://mmtf.rcsb.org) is the entry point to all documentation and software, including the MMTF specification, links to GitHub repositories of the MMTF APIs (Java, JavaScript, Python, C, and C++), and API descriptions. The versioned specification and all software libraries are available under either an Apache 2 or MIT license. A description how to download MMTF files is also available. A “Try it” feature demonstrates the transfer and parsing performance of MMTF-JavaScript in a web browser, and a “See it in Action” page demonstrates the fast data transfer, parsing, and rendering in NGL viewer [14].

Due to simple API, user-friendly specification and licensing model, the format has already been incorporated into several protein analysis tools and 3D structure visualization tools (Table 4).

**Table 4.** Applications that support the MMTF file format.

| Application | Link | Reference |
| --- | --- | --- |
| 3DMol.js | http://3dmol.csb.pitt.edu/ | [16] |
| BioJava | http://biojava.org/ | [15] |
| BioPython | http://biopython.org/ | [13] |
| ICM | http://www.molsoft.com/icm_browser.html |  |
| iCn3D | https://www.ncbi.nlm.nih.gov/Structure/icn3d/icn3d.html | [17] |
| JSmol/Jmol | http://www.jmol.org/ |  |
| MDAnalysis | http://www.mdanalysis.org/ |  |
| NGL | https://github.com/arose/ngl | [14] |
| PyMol | https://www.pymol.org/ |  |

We envisage the above advances will have a major impact in two areas of structural bioinformatics (Fig 12).

**Fig 12.**
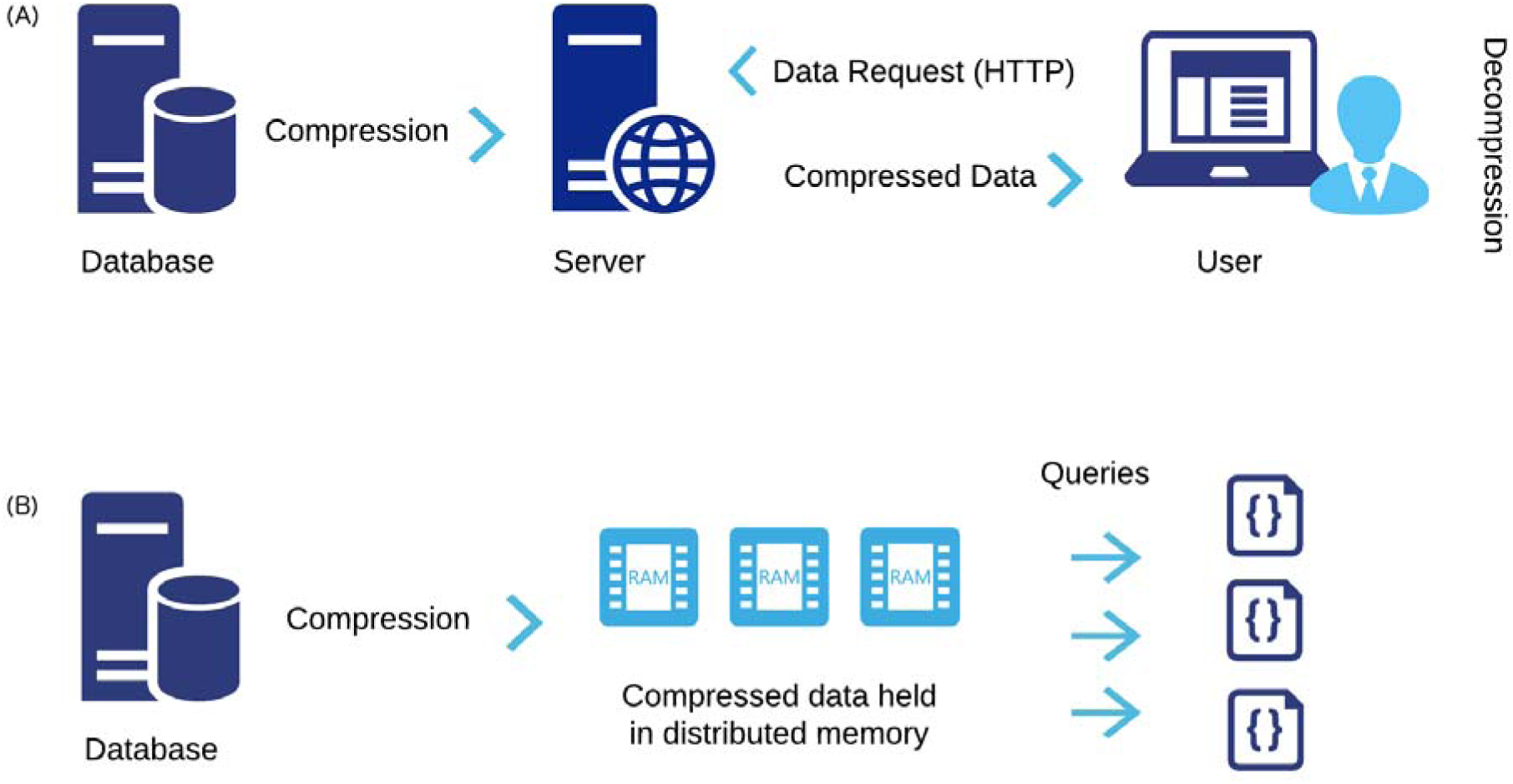
Main applications of the MMTF file format. (A) MMTF enables fast transfer, parsing, and low client side overhead for high-performance visualization in web-based viewers and in particular mobile devices. (B) MMTF can be represented in “Big Data” formats and the small size enables high-performance, in-memory analysis and calculations of the entire PDB archive using Big Data frameworks for parallel processing.

The first key area of impact is visualization of macromolecular structures, in particular when used on mobile device or in a web browser. MMTF enables low bandwidth file transfer, low client-side memory consumption, and fast parsing of PDB structures. For example, the 3D visualization on the RCSB PDB website is powered by MMTF [18], using the MMTF-full representation for entries < 10,000 residues and the MMTF-reduced representation for larger entries. Using the NGL viewer [14] and MMTF, the currently largest structure in the PDB, the HIV viral capsid (PDB ID 3J3Q) [2], can now be visualized on a mobile device (Fig 1A).

Second, by greatly increasing file-parsing speed, a rapid analysis of the entire PDB archive can be carried out. As an example, we have used the MMTF format to rapidly mine the PDB for interatomic distance distributions. Coupled with the use of an efficient geometric hashing algorithm in BioJava, the distances between all C-alpha carbons can be calculated in minutes. Parsing of the text-based mmCIF format alone would take several hours. Using a Hadoop Sequence file with MMTF records enables the scalable analysis of the PDB using standard distributed parallel processing frameworks. Further work is ongoing to demonstrate the use of MMTF with Big Data frameworks.

MMTF is an open source project and we welcome additions and new applications that use the new technology. As an example, the MMTF-C and MMTF-C++ decoders were developed in collaboration with community members.

## Acknowledgements

We thank Robert Hanson, Thomas Holder, and David Koes for their feedback on the MMTF specification and API. We thank Thomas Holder, Julien Ferté, Gazal Kalyan for developing the MMTF-C decoding library and Gerardo Tauriello, Stefan Bienert, Gabriel Studer, and Andrew Waterhouse for developing the MMTF-C++ decoding library. Robert Hanson provided efficient Java code for decoding of MessagePack. We also thank all users who helped with MMTF file transfer benchmarks worldwide, and Shih-Cheng Huang for performing the BioPython benchmarks. We thank Ezra Peisach for help with data validation, and Cole Christie and Chris Randle for setting up the weekly update process and data download for MMTF files.

